# YAP/TAZ mechanotransduction is a fast cellular response to subnuclear adhesions revealed by mechanically dynamic substrates

**DOI:** 10.1101/2025.06.04.656127

**Authors:** Alessandro Gandin, Veronica Torresan, Margherita Pelosin, Giada Vanni, Rebecca Busetto, Anna Citron, Ambela Suli, Tito Panciera, Francesca Zanconato, Stefano Piccolo, Giovanna Brusatin

## Abstract

Mechanotransduction, the conversion of mechanical cues into biochemical signals, is a cardinal regulator of myriad cell behaviors. However, the precise timing by which mechanical signals are sensed and relayed to the nucleus remains poorly understood. To address this gap, we developed dynamically degradable polyacrylamide (DPAA) hydrogels that allow in situ modulation of substrate stiffness while maintaining continuous cell adhesion, thereby mimicking physiological mechanical transitions. Using these tools, we discovered that YAP/TAZ, key mechanosensitive transcription factors, respond rapidly to changes in substrate stiffness primarily through changes in subnuclear adhesions. As substrate softening reaches the low kPa threshold (<2kPa), subnuclear adhesions and their associated ventral actin fibers disassemble within minutes, concurrently causing cytoplasmic relocalization of YAP/TAZ. In cells experiencing such a continuum of mechanical states, peripheral focal adhesions remained surprisingly stable while undergoing continuous remodeling, suggesting a more resilient and adaptable mechanism of force transmission than previously modeled. Our time-resolved results indicate that subnuclear adhesions act as rapid mechanotransduction nodes, leading to YAP/TAZ-mediated nuclear responses independently of peripheral adhesion dynamics. This enhanced understanding of YAP/TAZ regulation should advance the field of mechanobiology and offers potential insights for manipulating cellular responses in regenerative medicine and cancer therapy.

## INTRODUCTION

Mechanotransduction - broadly defined as the process by which cells convert mechanical cues from their environment into biochemical signals - is a fundamental driver of cell behavior, influencing everything from migration and differentiation to tissue homeostasis and disease progression^1–3^. Despite significant advances in understanding the molecular players involved in mechanotransduction, a persistent “black-box” remains around the precise timing by which mechanical signals are sensed at the cell surface and relayed to the nucleus. Dissecting the timing of nuclear mechanotransduction is crucial for multiple reasons. Mechanotransduction involves a complex network of interconnected events, and understanding the temporal order in which distinct cellular structures process this information helps clarify their hierarchy, cause-and-effect relationships, and primary pathways leading to nuclear effectors^4,5^. This would also allow differentiation between primary conduits and reinforcing or parallel events.

Addressing this timing gap is particularly challenging for at least two main reasons. First, defining the start and end of a given mechanical stimulus is rarely straightforward. In typical mechanobiology setups, cells are passaged, trypsinized and placed in suspension, and then plated on substrates of different stiffnesses or confined cell adhesive areas^2,6–11^. These setups abruptly switch cells from one mechanical environment to suspension and then to yet another mechanical environment, and this has so far limited the exploration of how cells adapt to dynamic mechanical transitions. This experimental set up also contrasts sharply with in vivo conditions, where cells remain continuously attached to the extracellular matrix or basement membrane and experience gradual or sustained changes in force. For example, in situ changes in extracellular mechanics are known to drive cellular reprogramming in the context of regeneration and cancer^1,12–17^, obviously leaving cells well adherent to their substrate.

A second caveat in studying the timing of mechanotransduction relate to the choice of its read out. For example, many studies rely on phenotypic outcomes such as growth or differentiation although these are slow biological processes, occurring over days and integrating multiple signaling pathways^18,19^. Notably, such feedback includes the fact that cells actively respond to mechanical cues by modulating ECM composition and mechanics (inside-out signaling^20^), an often-overlooked element of indetermination in mechanobiology assays. Irrespectively, long-term responses necessarily obscure the immediate events that initiate mechanosignaling and thus make it difficult to pinpoint exactly when the nucleus receives the mechanical message.

Together, the above challenges underscore the need for careful experimental design and innovative analytical tools recapitulating more physiologically-relevant transitions and more direct readouts of mechanical signaling. Only few pioneering efforts have recently initiated this endeavor by incorporating the temporal dimension into stiffness-controlled culture substrates. For instance, *in-situ* stiffening or softening hydrogels have been developed using light assisted modifications of the crosslinking degree of the adhesive substrate^21–23^. However, preparing these photo-tunable substrates has drawbacks: changes in modulus do not occur seamlessly, but rather as discrete step changes imposed by light-assisted softening or defined by the experimenter. These set ups also prevented so far the ability of live-image cells during their continuous adaptation to a mechanically dynamic substrate.

With this background in mind, we aimed to overcome existing limitations by developing new biomaterials that allow to visualize dynamic mechanosensing in living cells. Our findings revealed that mechanosignaling occurs through at least two distinct steps, a first one closely associated to nuclear vs. cytoplasmic localization of the YAP/TAZ transducers^1^, and a later step, primarily concerned with the restructuring of the cellular adhesive architecture. We further revealed the central role of subnuclear adhesions and basal actin fibers as the onset of YAP/TAZ mechanotransduction.

## RESULTS

### Synthesis of in-situ degradable adhesive hydrogels

To investigate the timing of cellular mechanotransduction from the extracellular matrix (ECM) to the nucleus we developed a cell-compatible hydrogel system that permits rapid, in situ modulation of substrate stiffness. This design allows cells to remain continuously attached to their substrate as the latter transits between a continuum of mechanical states. By maintaining cell-substrate contacts throughout these mechanical perturbations, we can more faithfully capture the initial events and subsequent adaptations that define a cell’s response to altered ECM rigidity. This set-up should more closely recapitulate physiological conditions while avoiding artifacts associated with detachment and re-plating.

For the hydrogel composition, we selected polyacrylamide (PAA) substrates functionalized with RGD adhesive peptides. PAA’s chemical tunability, well-established fabrication protocols and widespread popularity for mechanobiology studies, make it an ideal choice for precise control over mechanical properties^2,24^, while the presence of RGD ensures reliable, integrin-mediated cell adhesion. To impart dynamic control over stiffness, we envisioned an innovative procedure: we incorporated during synthesis disulfide (S–S) crosslinkers (**Figure 1a**), which can be cleaved to thiols (SH) through reduction^25–27^. For this reduction step, we opted to use glutathione (GSH), a naturally occurring molecule that is maintained at high millimolar concentrations (up to 10 mM)^28^ inside cells to prevent oxidative damage^29^, yet exists at negligible levels extracellularly. Because GSH is cell-impermeable, adding even a low-concentration spike of it to the culture medium provides a highly effective “switch” to soften the disulfide-crosslinked PAA – hereafter referred to as dynamic (DPAA) hydrogels.

**Figure 1:**
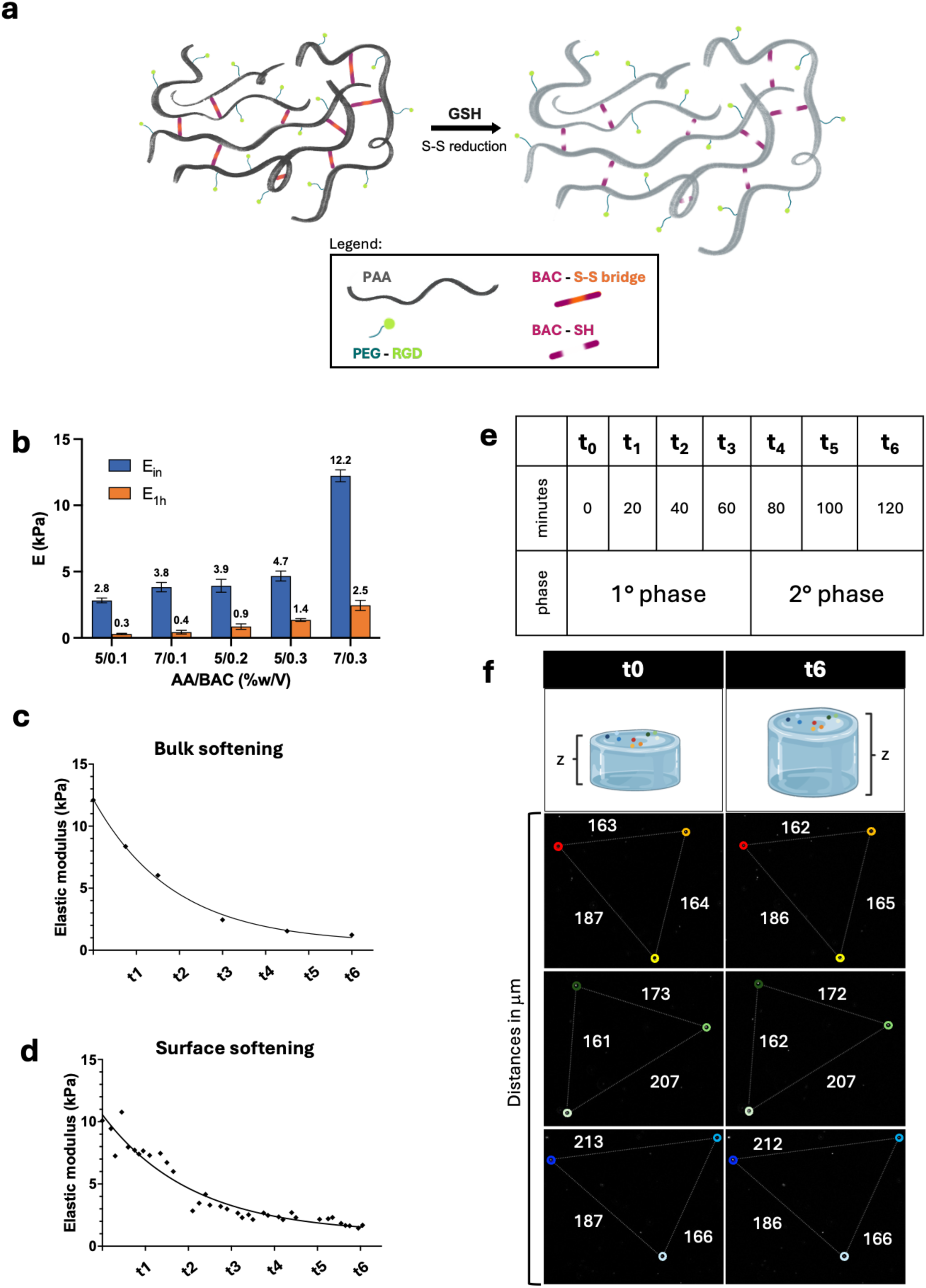
a) Scheme of the DPAA gel and of its softening by degradation of S-S bridges with the GSH addition. For the synthesis, Acrylamide (AA), bisacryloyl-cystamine (BAC) and a PEG monomer conjugated with a RGD containing peptide are copolymerized with a radical reaction. Different ratio between AA and BAC are used to achieve tunable initial stiffness in the range of about 3-12kPa (**Table S2**). All chemical compounds used are reported in **Table S1**. b) Initial and final stiffnesses of gels with different formulations degraded for 1h with 1mM of GSH (see details in **Table S2**). (c) Bulk stiffness measured at different time points during softening with 1mM GSH of a gel with initial stiffness of about 12kPa (see details in **Table S3**). (d) Surface stiffness measured with AFM at different time points during softening of a gel with initial stiffness of about 12 kPa. (e) Time window – timepoints correlation. Biphasic behaviour shows a first phase corresponding to softening between t_0_ – t_3_ and a second phase extending between t_4_ - t_6_. Timing may vary by up to 10 minutes earlier or later depending on the individual cell examined (f) Lateral swelling behavior investigated by confocal microscopy on gels embedding fluorescent particles (diameter 0.5μm). Lateral swelling is measured monitoring the variation of their distance during 2h softening (1mM GSH)

DPAA were synthesized by radical copolymerization of various formulations of acrylamide (AA) and bisacryloyl-cystamine (BAC) in water (**Figure 1b and Table S1**). For RGD functionalization, we first conjugated a maleimide-PEG-acrylate monomer with a short RGD peptide, creating an acrylate-modified adhesive peptide (AC-PEG-RGD). Then, AA and BAC were co-polymerized with AC-PEG-RGD, each at fixed concentration of 3mM for all gels (see details in **Tables S2**). The introduction of a PEG chain was to overcome a known limitation of PAA hydrogels functionalized with adhesive peptides, that is the presence of PAA “brushes” on the gel surface, which inhibit the availability of adhesive sequence to cellular integrin receptors.^10,30^

To optimize the dynamic range of DPAA stiffness (i.e., the modulus values before and after GSH-induced softening), we synthesized hydrogels at various initial rigidity levels by varying both monomer and crosslinker concentrations (**Table S2**). After 1 mM GSH was added for one hour, we measured the modulus changes via micropipette aspiration (**Table S3**). Results showed that DPAA gels could initially reach stiffness values of approximately 3 to 12 kPa, which then softened by up to tenfold – for instance, dropping from 3 kPa to 0.3 kPa – as such spanning the entire spectrum of physiologically relevant ECM rigidities (**Figure 1b and Table S2**).

Next, we investigated the temporal dynamics of hydrogel softening. All DPAA gels exhibit a significant decrease in stiffness within the first hour of GSH treatment, regardless of their initial rigidity (**Figure 1b**). After this initial phase, the gels reach a plateau of residual stiffness that remains stable even upon extending the GSH exposure to four hours (**Figure S1**). This plateau is likely attributable to PAA chain entanglements in the gel network rather than GSH depletion via oxidation, as replenishing GSH does not further reduce stiffness. We also measured mechanical changes at the surface level as this is what cells experience. Atomic force microscopy (AFM) measurements on the adhesive surface confirmed that DPAA hydrogels undergo a comparable modulus shift and timing of softening when probed directly at their cell-contacting interface (**Figure 1d**).

Next, we aimed to exclude the possibility that reduction of hydrogel crosslinking could affect mechanosignaling indirectly by changing the superficial density of adhesive ligands due to lateral swelling.^10^ For this, swelling behavior was investigated by particle tracking during 2h after GSH addition. As shown in **Figure 1f**, substrates did not exhibit substantial changes in their lateral dimensions during 2h of softening and swell only along the z axis (**Figure S2)**, confirming that softening does not alter the density of adhesive ligands.

### Live imaging of adherent cells on softening gels reveals their gradual adaptation to substrate stiffness

The most mechanistically understood step in mechanosignaling occurs at the cell surface, that is, integrin clustering into focal adhesions^31,32^. The latter sense and transmit mechanical forces from the cell microenvironment to the actin cytoskeleton, with actin stress fibers and other cytoskeletal elements, including microtubules and intermediate filaments, relaying forces throughout the cell. A key outcome of this force transduction is the nuclear translocation of mechanosensitive transcription factors such as YAP/TAZ^1,2^, though the precise molecular mechanisms by which F-actin exerts control over their activity remain incompletely understood. An emerging concept is that the nuclear envelope functions as a mechanosensitive organelle^33–35^, yet the causal links among subcellular structures, the temporal phases of mechanotransduction, and the specific molecular players at each step of force propagation remain to be fully elucidated.

Leveraging the dynamic mechanical properties of DPAA hydrogels, we conducted live-cell imaging experiments to observe how cells adapt in real time to changes in substrate stiffness. Specifically, we tracked the nuclei, actin cytoskeleton, and focal adhesions of WI38 human fibroblasts grown on gels initially measured at ∼12 kPa (**Movie 1 and 2**, **Figure 2a** and **2c**). To capture real-time shifts in F-actin structure during the softening process, we used Fast-Act, a minimally perturbing live probe that allows natural actin assembly and disassembly.

**Figure 2:**
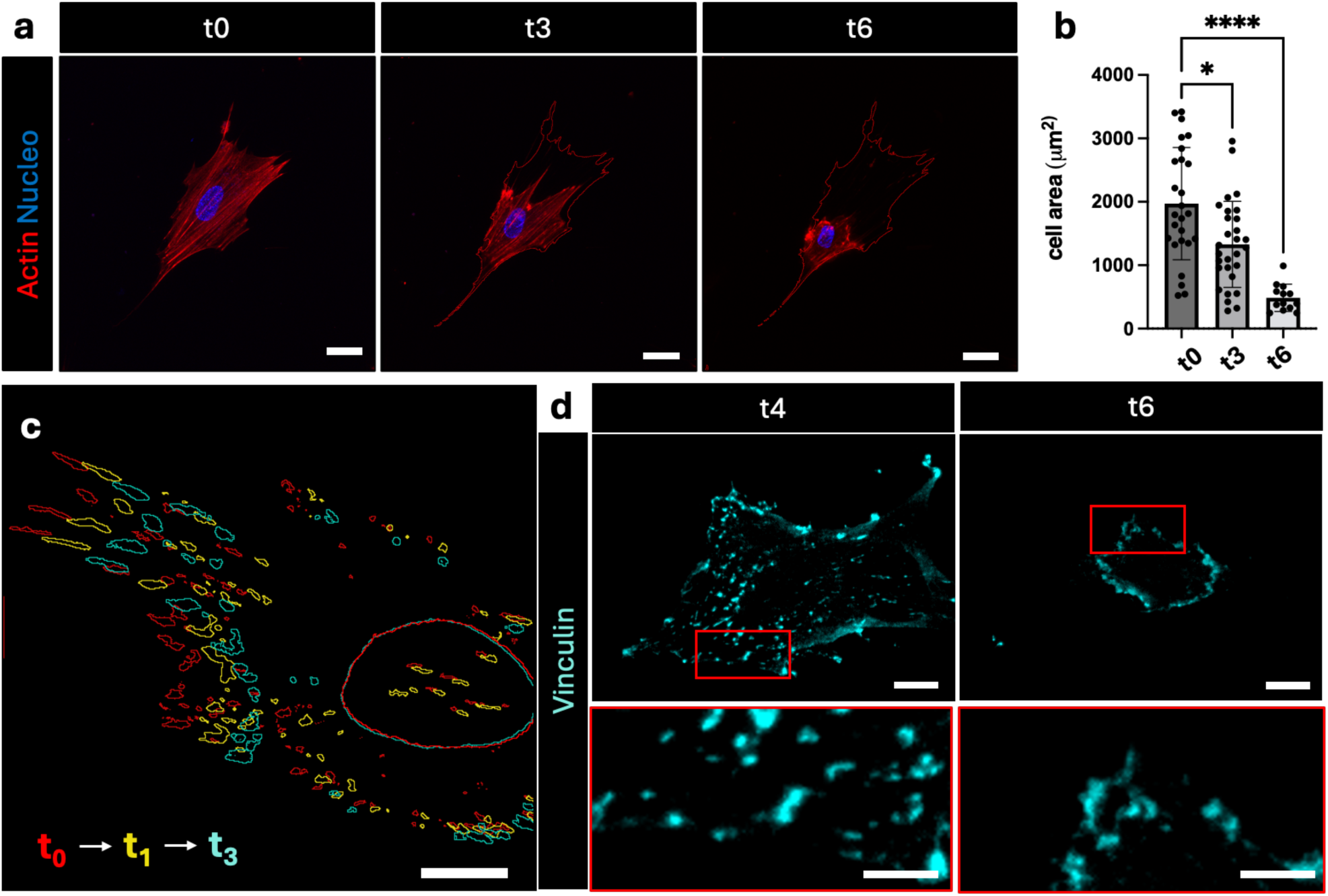
**a)** Pictures taken during live imaging of nuclei and actin cytoskeletons of WI38 during gel softening up to t_6_. Live imaging probe (Fast-act) is used to minimally perturb F-actin dynamics. (full movies in **Movie 1).** Scale bar 20 μm. b) Cell area temporal evolution during softening up to t_6_. Data are reported as means and standard deviations. Statistical significance is evaluated with one-way ANOVA with Welch correction (p values: t_0_-t_3_: 0.0131; t_0_-t_6_: <0.0001). Cell area and nuclear projected area are measured on fixed samples stained with fluorescent phalloidin and Hoechst nuclear dye. c) Live imaging of WI38 cells infected with Vinculin Venus (full movie in **Movie 2**). Cells are seeded on gels of about 12kPa as initial stiffness, allowed to adhere to the hydrogel surface overnight and imaged during gel softening (1mM GSH). Nuclear outline is obtained by Hoechst staining. Scale bar 10 μm. d) Representative images of the WI38 cells seeded on 12kPa DPAA gel and imaged during the second phase (t_4_ - t_6_). Cells are fixed at t_4_ and t_6_ and stained for vinculin. Scale bar 10 μm for low magnification images and 5 μm for zoom-in

We made several interesting observations. At t_0_, the cells were stretched and showed abundant stress fibers aligned with their main axis, indicative of strong actomyosin contractility (**Figure 2a left panel**). After glutathione pulse, cells progressively shrink from their peripheral projections (**Figure 2b**), shortening the underlying stress fibers, but retaining their initial orientation and axial organization. Moreover, we noticed a biphasic response in focal adhesions and peripheral stress fibers. During the first phase (**t_1_ – t_3_ of Figure 1c**), corresponding to substrate softening from 12 kPa to 2 kPa, peripheral adhesions do not disappear. Instead, they gradually retract toward the cell center while decreasing in length and thickness. These transitions occur over a timeframe ranging between 25 to 60 minutes, depending on the individual cell examined.

In order to more precisely assess changes in cell-substrate engagement we monitored the dynamics of mature focal adhesions by using a fluorescent vinculin reporter. Live imaging of cells transfected with venus-tagged vinculin (**Movie 2**), indeed revealed that most mature focal adhesions remain intact during this first phase of softening; interestingly, rather than disassembling, they exhibit a continuous, crawling-like movement toward the cell center (**Figure 2c**). This suggests that pre-formed focal adhesions are self-organizing and self-preserving. Up to this stage, the nuclear projected area and nuclear positioning show much less dynamic behavior, with minimal shrinkage over the same timeframe.

The second phase occurs rapidly, (t_4_-t_6_ of **Figure 1d**) from about 60 to 120 minutes during which substrate stiffness drops from 2.5 kPa to 1.5 kPa and this is accompanied by shortened peripheral adhesions suddenly and collectively disassembling (**Figure 2d**) and concomitant dramatic cell shrinkage (**Figure 2a right panel**).

### YAP/TAZ is an immediate response to cellular mechanosensing

Having established a biphasic response in cellular adaptation to mechanically dynamic substrates, we next investigated which of the observed adhesive, cytoskeletal and cell shape changes is most directly linked to mechanically controlled transcriptional responses. To do this, we matched YAP/TAZ localization with changes in cytoskeletal organization, cell projected area and nuclear shape. Specifically, we performed a series of quantitative analyses comparing these parameters at time 0 (t_0_), at the end of the first and second phases of mechanosignaling (i.e., at t_3_ and t_6_, respectively). For this, we used DAPI to monitor nuclear shape and phalloidin staining to gauge the cell projected area. We found that in cell exposed to our dynamic substrate, YAP/TAZ are nuclear at the initial stiffness of 12 kPa, as expected. We found that at t_3_, cells already exhibited cytoplasmic YAP/TAZ without displaying any overt alterations in classic parameters of cellular mechanorespose such as cell shape, nuclear projected area and nuclear aspect ratio, at least at the end of the first phase (**Figure 3**). Indeed, alterations in nuclear aspect ratio, and overall cell rounding are later events that do not initially drive YAP/TAZ mechano-regulation (**Figure 3c** and **3d**) and that occur from t_4_ to t_6_.

**Figure 3:**
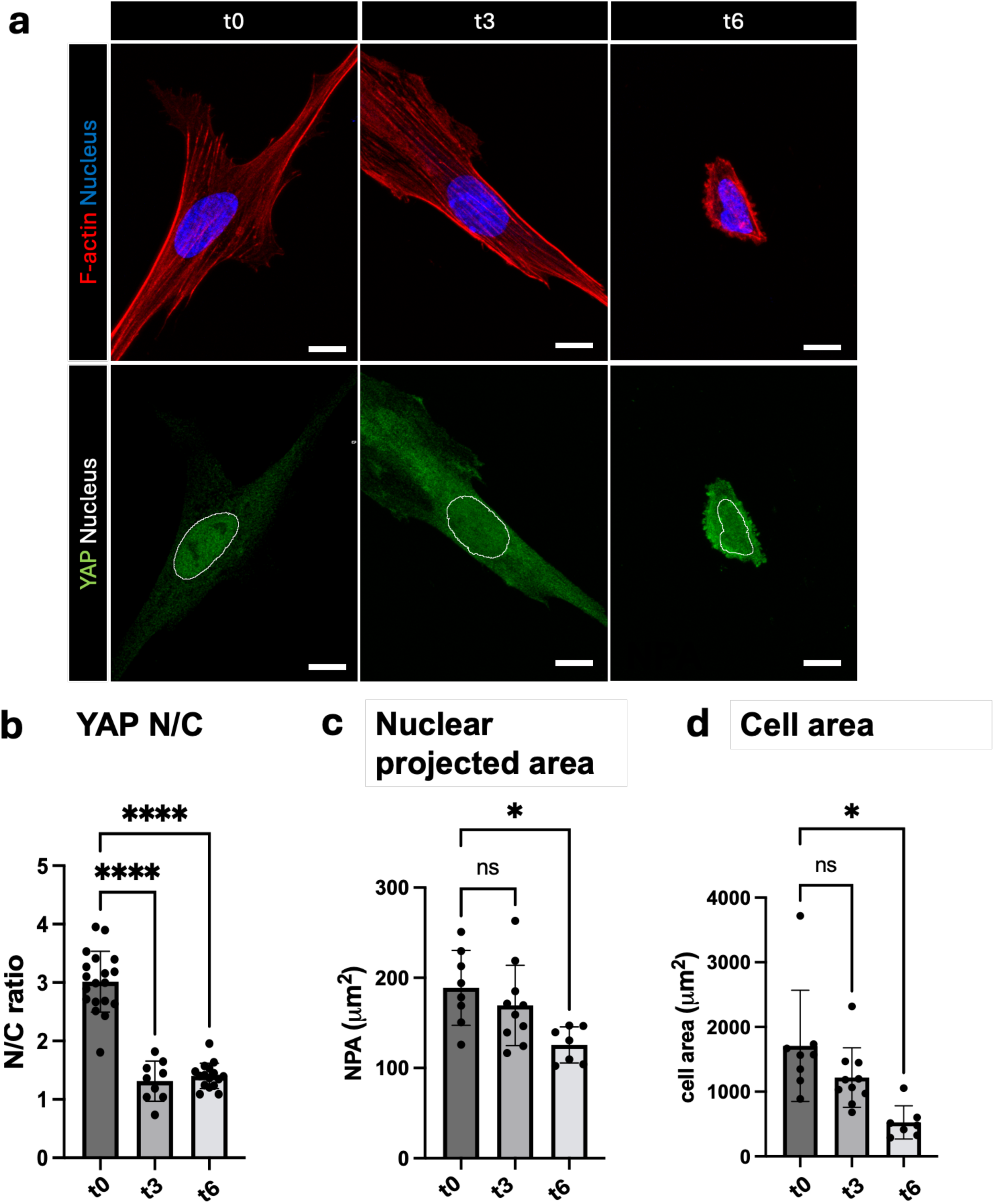
a) Representative immunofluorescence images of WI38 seeded on cells seeded on 12kPa DPAA gel at different time points during gradual softening of dynamic gels. From the staining are visible: nuclei (in blue), YAP/TAZ (in green) and F-actin (in purple). F-actin was stained with fluorescently labeled phalloidin to serve as cell shape reference. All cells were seeded for 24h before analysis and then fixed and stained at different time points (t_0_, t_3_, t_6_), to which correspond different stiffnesses as indicated in Figure 1c. Scale bar 10 μm b) YAP/TAZ quantification measured by the Nuclear to Cytoplasmic (N/C) ratio of YAP/TAZ subcellular localization. (p values t_0_-t_3_: <0.0001; t_0_-t_6_: <0.0001) c) Nuclear Projected Area (NPA) analyzed from Hoechst fluorescent signal (p values t_0_-t_3_: 0.7120; t_0_-t_6_: 0.0095) d) Cell area analyzed from the fluorescently tagged phallodin signal (p values t_0_-t_3_: 0.4179; t_0_-t_6_:0.0168). Data in b, c and d are reported as means and standard deviations. Statistical significance is determined with one-way ANOVA with Welch correction

Next, we examined other structural features, focusing on peripheral focal adhesion composition and length as an inside-out proxy of cellular traction forces. To explore focal adhesion clustering, we assessed the clustering of individual integrins, specifically α_5_β_1_ and α_v_β_3_, that are main RGD-binding integrins (**Figure 4a** and **4b**). Similar to our findings for vinculin, the density of these adhesion sites at the cell periphery remained unchanged. Overall, these results suggest that peripheral adhesions are unlikely to serve as immediate regulators of YAP/TAZ activity.

**Figure 4:**
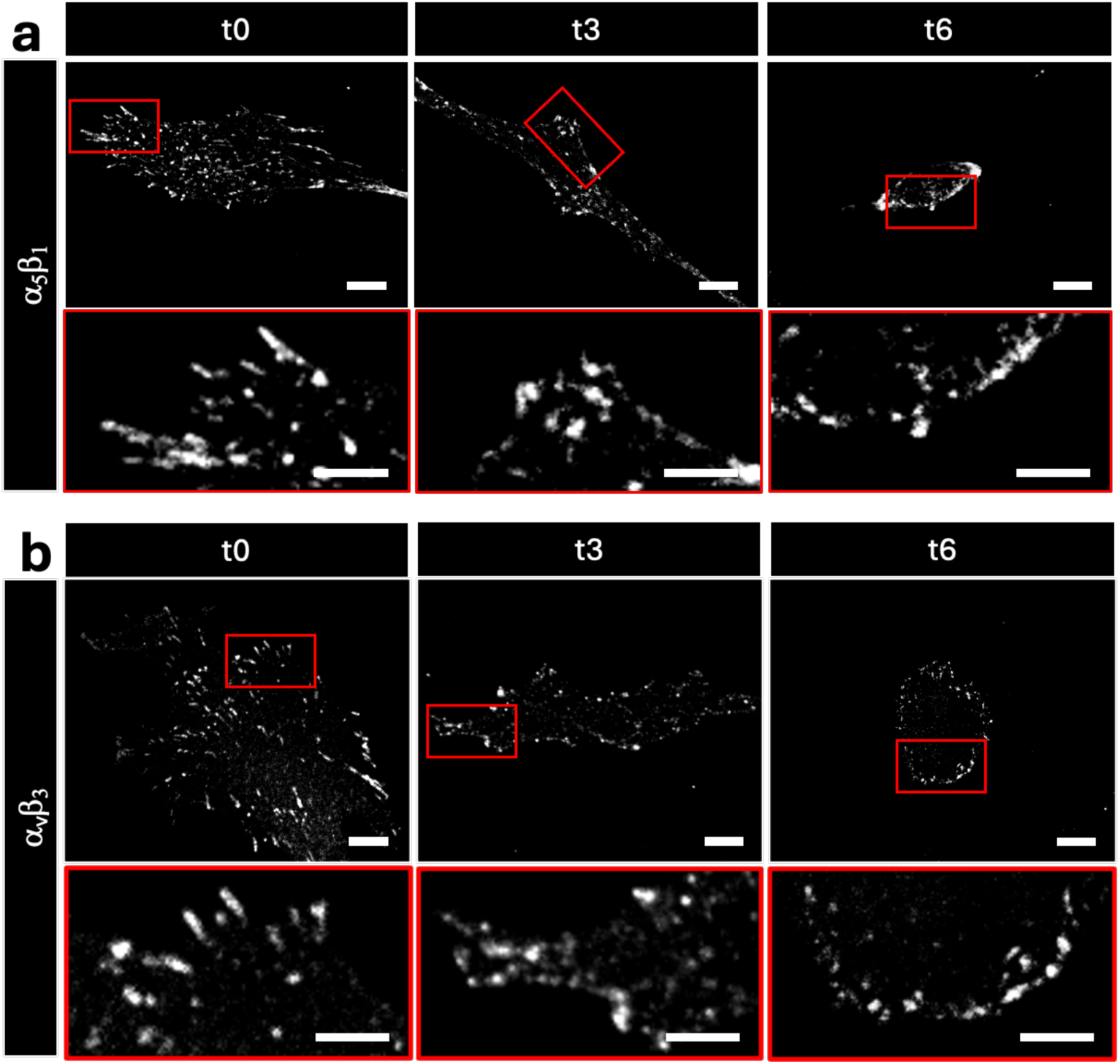
Immunofluorescence images of WI38 seeded on DPAA gels with about 12kPa of initial stiffness at different time points during gradual softening. From the staining are visible a_5_b_1_ integrins (a) and a_v_b_3_ integrin (b). All cells were seeded 24h before analysis, then fixed and stained at different time points (t_0_, t_3_, t_6_) to which correspond different stiffnesses as indicated in Figure 1c. Scale bars 10 μm and 5 μm for zoom-in

We started this investigation aiming to correlate how dynamic changes in substrate stiffness affect cellular mechanotransduction. However, the above results raised an intriguing conundrum: despite detecting expected changes in YAP/TAZ localization, these occurred before overt modulation of a number of parameters (adhesive, cytoskeletal and nuclear) that have been previously linked to mechanoregulation at least in steady state conditions. This prompted us to more closely investigate the spatial, intracellular location, of adhesive subpools. In particular, by live imaging of vinculin dynamics, our attention was attracted on the dynamic of another subtype of adhesive element, that is subnuclear adhesions and their associated ventral stress fibers. Specifically, we noted that, differently from peripheral adhesions, changes in these adhesive sites occurs in the first phase of mechanosignaling and are temporally proximal or concomitant to YAP/TAZ nuclear exclusion. To better visualize and confirm this results we used α5β1 and αvβ3 signals and also an independent marker of focal adhesions, i.e., paxillin (**Figure 5 and Figure S3**). We found that, in stark contrast with peripheral adhesions, subnuclear adhesions are already overtly disassembled at t_3_. This temporal analysis raised the intriguing possibility that YAP/TAZ regulation is primarily driven by tethering of F-actin to subnuclear focal adhesions, whose “first to go” disassembly configures a fast response to changes in substrate physical properties.

**Figure 5:**
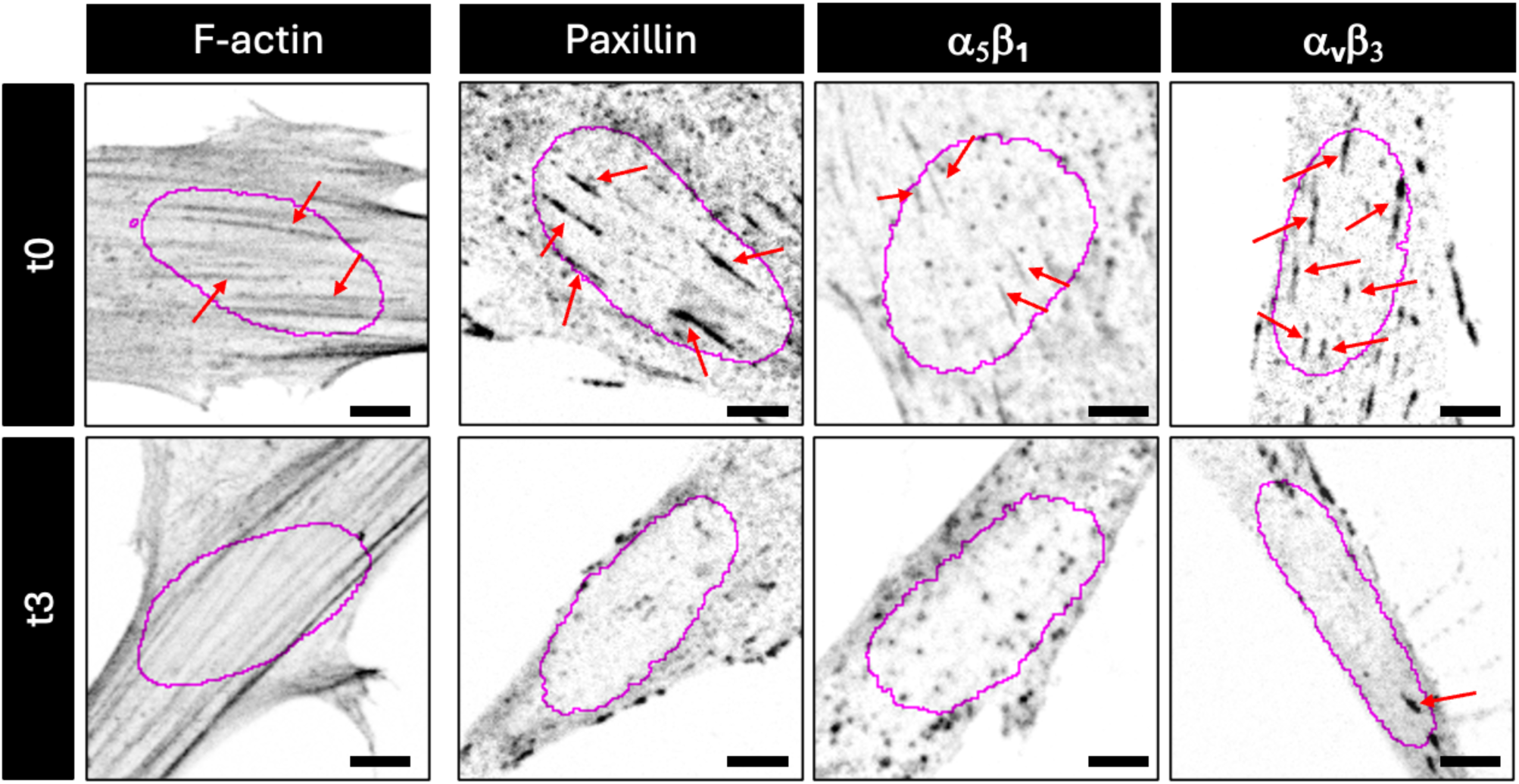
Representative immunofluorescence images of cells on degradable hydrogel at 12 kPa (t_0_) and 2.5 kPa (t_3_). Subnuclear basal actin fibers and cellular adhesions (paxillin, a_5_b_1_ or a_v_b_3_) were imaged through confocal microscopy. Purple circles represent the outline of the nuclear projected area. Red arrows indicate the main subnuclear adhesions which colocalize with the terminations of thick actin bundles in cells perceiving 12 kPa. Adhesions disassembly associates with a remodeling of the basal actin fibers in thinner and longer filaments. Scale bar 5 μm

To corroborate the above conclusions, we directly quantified YAP localization and subnuclear vs. peripheral adhesions in more time-resolved window, assessing these values around t_3_ with a 10’ resolution within the same cell. Strikingly, loss of YAP/TAZ nuclear localization occurs suddenly in just 10 minutes preceding t_3_ and this occurs concomitantly to disappearance of subnuclear adhesions (**Figure 6a** and **b**). Of note, the latter is also accompanied by loss of the specific subpool of F-actin stress fibers that tether the nucleus to the basal/ventral adhesive side of the cell. From this point onwards, YAP/TAZ localization and nuclear tethering to basal/ventral stress fibres do not further change (**Figure 6a, right panels**). The results suggest that YAP/TAZ nuclear exit occurs concomitantly to the loss of nuclear tethering to subnuclear stress fibers that are in turn anchored to subnuclear adhesive sites.

**Figure 6:**
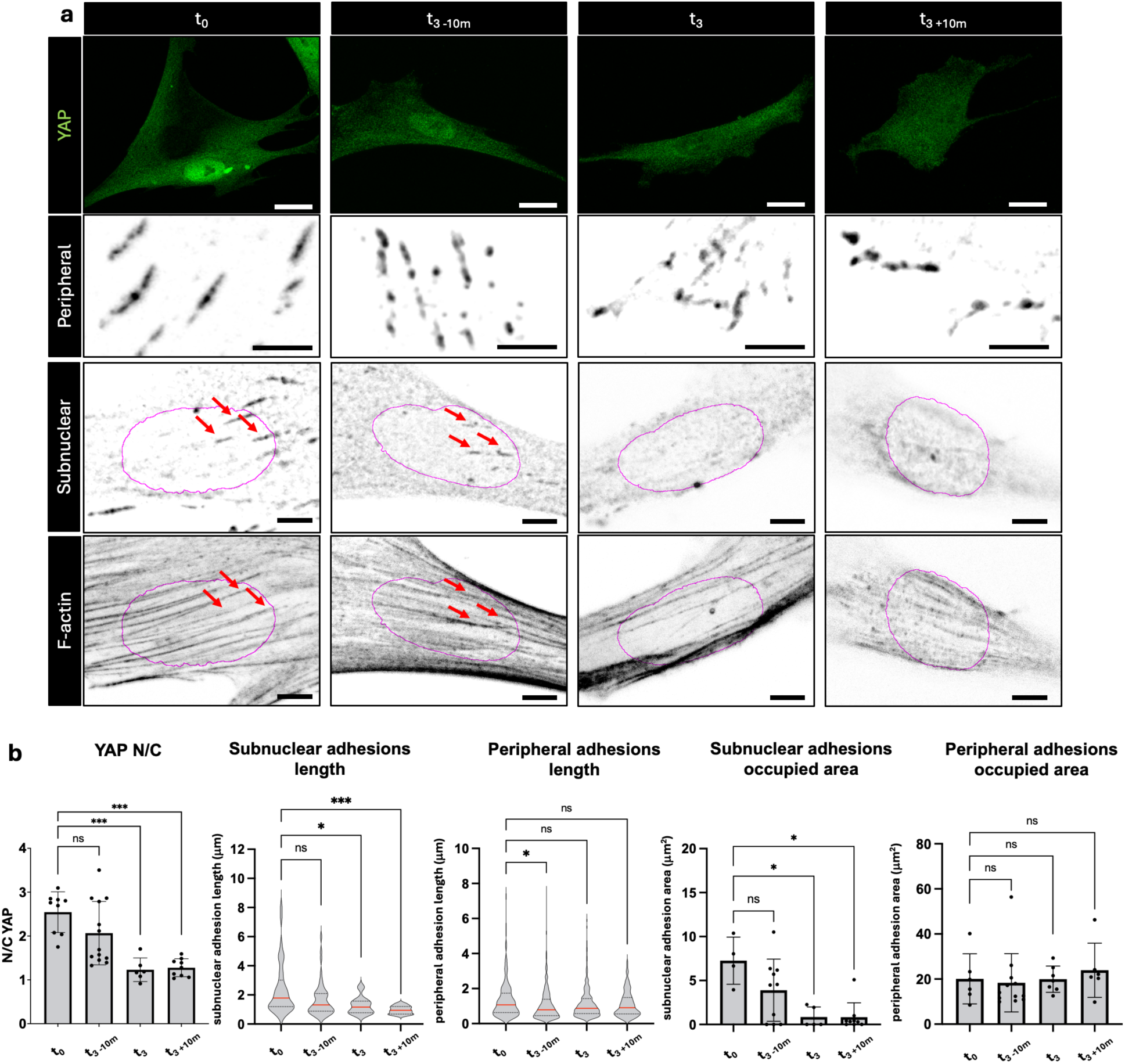
Temporal evolution of YAP/TAZ localization and peripheral and subnuclear adhesions quantified in short interval timepoints by YAP/TAZ and paxillin immunofluorescence. a) Representative images of YAP/TAZ, peripheral and subnuclear paxillin, and basal actin signals for each timepoint. Purple circles indicate the outline of the nuclear projected area obtained by Hoechst counterstain. Scale bars 20 μm for YAP/TAZ, 5 μm for adhesions and F-actin. b) Quantification of YAP/TAZ localization, length and area of subnuclear and peripheral adhesions. Statistical significance for YAP localization, subnuclear adhesion length and subnuclear adhesion area were evaluated with one-way ANOVA with Welch correction. p values (for YAP t_0_ – t_3-10_: 0.4871; t_0_-t_3_: 0.0001; t_0_-t_3+10_:0.0001, for subnuclear adhesions length t_0_-t_3-10_:0.2097; t_0_-t_3_:0.0258; t_3_-t_3+10_:0.0002 for peripheral adhesions length: t_0_ – t_3-10_: 0.0148; t0-t3: 0.1632; t_0_-t_3+10_:0.0811, for nuclear adhesions area t_0_-t_3-10_: 0.3822; t_0_-t_3_: 0.0467; t_0_-t_3+10_: 0.0475, for peripheral adhesions area t_0_-t_3-10_: 0.9997; t_0_-t_3_: >0.999; t_0_-t_3+10_: 0.9907)

All in all, our dynamic hydrogels reveal that the timing of YAP/TAZ mechano-regulation is, at least in its incipit, exquisitely dependent on the dynamics of subnuclear adhesions in response to changes in substrates rigidity.

## DISCUSSION

The present results unveil the timing by which mechanical information translates into nuclear events, an aspect of cell signaling that was so far enigmatic. Mechanotransduction, by its very nature, entails a myriad of interconnected events across different scales, from integrin clustering at focal adhesions to complex cytoskeletal reorganizations and nuclear envelope deformations. Navigating through this complexity can be daunting. Here, we fixed our compass on one dominant event of mechanotransduction: the nuclear accrual of YAP/TAZ, by which mechanics ultimately governs gene expression. By zeroing in on the timing of this fundamental signaling step, we were able to pinpoint the prime relationships and functional interdependencies between the earliest subcellular that trigger downstream transcriptional responses and YAP/TAZ activation. Equally important, we distinguished these early events from subsequent stages of cell reorganization that might be secondary, independent or reinforcing the core nuclear mechanosignaling pathway. This refined temporal map of mechanotransduction offers new insights into how cells initially sense and interpret mechanical cues, setting the stage for targeted approaches to manipulate early mechanosensitive processes in various physiological and disease contexts.

Our work contains several elements of interest. The first is methodological. In order to address the spatiotemporal challenges of mechanotransduction, we established a dynamic polyacrylamide hydrogel platform that can be rapidly softened in situ. These hydrogel formulations do not require sophisticated implementations and can be readily adopted in any cell biology laboratory. This platform allows us to overcome a hitherto major limitation in mechanobiology research, namely, the inability to continuously modulate substrate stiffness while maintaining persistent cell-substrate contacts. Indeed, traditional approaches often rely on abrupt switching - detaching cells and re-plating them on substrates of different rigidity - or on phototunable gels that undergo discrete, step-like changes ^21–23^. In contrast, the dynamic polyacrylamide hydrogels described here provide a gradual and controlled transition in stiffness, ensuring that cells remain attached to their underlying ECM and can be monitored continuously throughout the softening process.

Leveraging our dynamic hydrogel platform, we could carry out unprecedented live imaging experiments to monitor changes in F-actin organization and Focal Adhesions occurring during cellular adaptation to gradually decreasing substrate stiffness. Our observations revealed that preformed focal adhesions are simultaneously stable and highly dynamic: rather than collapsing in response to reduced force, these adhesions maintain cell–substrate attachment by shifting centripetally, thereby continuously adapting to the decreasing rigidity of the substrate. This behavior marks, at least in part, a step forward in our understanding of stiffness sensing. The latter has been so far mainly conceptualized accordingly to a “molecular clutch” model, largely inferred from static cellular studies in cells experiencing substrates of fixed degrees of stiffness^36,37^. According to the molecular clutch model, on softer ECM, substrate deformation absorbs part of the cell-generated force, reducing the rate of force transmission from integrins to F-actin. In this view, the rate of F-actin engagement at focal adhesion is slower than the integrin-ECM bond lifetime, ultimately causing bond dissociation and focal adhesion collapse. Our real-time analyses of cells transitioning from a stiff to a softer substrate revealed a more nuanced scenario as, in these dynamic conditions, focal adhesions only partially disassemble to continuously remodel themselves. They crawl backward on progressively softer substrates, in a binding continuum to nearby adhesive sites while preserving cell adhesion, contractility and structural integrity. This implies that when cells are initially adhered to a highly rigid extracellular matrix (ECM) and subsequently experience decreasing pulling forces, the molecular clutch remains engaged. We postulate that this is due to the fact that previously engaged focal adhesion remains as such for a time sufficiently long to allow a “treadmilling” turnover of new integrin molecules. In this view, as ECM pulling forces diminish and cell retracts, new integrins are continuously incorporated in focal adhesions binding to neighboring RGD adhesive sites, effectively substituting those that disengage at the periphery of the adhesion complex. This dynamic process ensures that force transmission is maintained in fluctuating mechanical environments, even as substrate stiffness decreases, highlighting the adaptability and resilience of the molecular clutch in supporting continuous mechanotransduction. At later time points, as softening continues, ECM rigidity reaches a critical threshold below which peripheral adhesions rapidly collapse, leading to increased cell rounding, that is the typical shape of cells plated on soft substrates in classic mechanobiology studies. In other words, by observing these processes in real-time, our dynamic hydrogel platform provides critical insights into the early stages of mechanosignaling, revealing how cells can finely tune their adhesive structures to maintain functionality in fluctuating mechanical environments.

A key finding involves the use of the nuclear versus cytoplasmic localization of YAP/TAZ to better understand both the timing of the cell’s mechanosensory response and its association with specific subcellular structures. Specifically, we discovered that the early disassembly of subnuclear adhesions is linked to the inhibition of YAP/TAZ. Our findings revealed that subnuclear adhesions – adhesive complexes located beneath the nucleus – disappear earlier during the softening process than peripheral adhesions. Notably, this timing coincides with the nuclear exit of YAP/TAZ. Through time-lapse imaging of subnuclear adhesion and YAP/TAZ localization within individual cells, we observed that cells retain intact, mature peripheral adhesions for the first phase of mechanosensing.

The rapid temporal pattern of subnuclear adhesion disassembly and YAP/TAZ inhibition indicates that subnuclear focal adhesions and their associated ventral actin fibers are the primary structures responsible for sensing and transmitting mechanical changes to the nucleus, and, through still undefined mechanisms, to cytoplasmic localization of YAP/TAZ. We can only postulate, at this point, that the early disassembly of these subnuclear adhesions may impact on one or more mechanical rheostats, either triggering inhibition of a nuclear YAP/TAZ anchoring factor or stabilization/activation of a cytoplasmic YAP/TAZ sink. Time-resolved investigations using DPAA gels should facilitate the identification of these mechanical sensors.

It is worth discussing that several factors could contribute to why subnuclear focal adhesions (FAs) might be particularly sensitive to ECM softening compared to peripheral adhesions. In one model, subnuclear FAs connect F-actin to the nucleus, allowing force transmission to nuclear envelope (including LINC complexes), and not to another FA as it occurs more peripherally. As non-alternative scenario being subnuclear actin cables shorter than peripheral ones, they may disassemble when subjected to comparatively lower mechanical forces when compared to the latter. One of the appeals of the latter model is that it may explain the behavior of cells forced to adhere to small adhesive islands: despite sensing a stiff ECM (at plastic rigidity), these “small” cells can only form short microfilaments and indeed display YAP/TAZ inhibition and phenotypic responses indistinguishable from those experienced on soft ECM. Irrespectively, the above combined factors may position subnuclear adhesions as key early responders to mechanical cues, enabling rapid modulation of YAP/TAZ localization before peripheral adhesions fully disengage during the mechanotransduction process.

In conclusion, the DPAA system provides a versatile platform for dissecting short-lived, early mechanotransduction events that traditional approaches often overlook. By preserving continuous cell-substrate interactions during substrate softening, we unveil a new dimension in the timing of mechanotransduction, suggesting that efforts to modulate subnuclear adhesion dynamics could be a potent strategy to control YAP/TAZ-driven cellular processes in development, regeneration, and disease.

## MATERIALS AND METHODS

### Non-adhesive glass preparation

Non-adhesive hydrophobic glass slides were used as platform for the polymerization of DPAA hydrogels. A 1 mm thick glass substrate was first washed with water to remove any residual grease or powder from the surface. Then, each side of the slide was submerged in a 3N sodium hydroxide (NaOH) solution for 15 minutes to clean and prepare the glass slide for silanization. After repeated washing with water, the slide was left to dry in an oven at 140°C for about 40 minutes. After surface activation with a plasma cleaner (Harricks Expanded Plasma Cleaner), the glass slide was silanized by covering the surface with PlusOne™ Repel-silane ES (Cytiva no. 17-1332-01). After 15 minutes, the slide was washed three times with pure ethanol and was left to dry in an oven at 80°C before use.

### Synthesis of DPAA hydrogels

To synthesize various formulations of dynamic polyacrylamide (DPAA) hydrogels, acrylamide (AA) solution (40 wt% in water, Sigma-Aldrich no. 01697), bisacryloyl-cystamine (BAC, Sigma-Aldrich no. A4929) solution (3 wt% in AA 40 wt%) and water were mixed to achieve the following compositions 5%AA 0.1%BAC or 7%AA 0.1%BAC or 5%AA 0.2%BAC or 5%AA 0.3%BAC or 7%AA 0.3%BAC. To introduce adhesive sites on the gel surface, the solution of AA and BAC was copolymerized with an acrylate-PEG-maleimide monomer (Lysan Bio, ACRL-PEG-MAL-5000) previously conjugated with amino-cysteine terminated synthetic peptide containing the adhesive sequence RGD, GRGDSPC. The conjugation reaction was carried out in water at room temperature: the RGD peptide and acrylate-PEG-maleimide were dissolved to a final molar ratio of 1:1 and mixed with the prepolymer solution at a fixed concentration of 3 mM for all formulations. Monomers solution was degassed for 15 min at 0.1 bar, to remove the dissolved oxygen which would inhibit the polymerization. Once degassed, the monomer solution was mixed with 1% v/v of 10 wt% ammonium persulfate (APS, Sigma-Aldrich 09913) solution and 0.1% v/v of tetramethylethylenediamine (TEMED, Sigma-Aldrich T7024). Once mixed, the solution was poured inside proper PDMS molds sealed with functionalized glass coverslips and it was left to polymerize for 20 minutes.

The PDMS molds and the glass coverslip dimensions are optimized according to the specific experiment: 250 μm thick PDMS ring and #1 18 mm round coverslips were used for fixed-cell imaging, 50 μm thick PDMS stripes and #0 25 mm round coverslips were used to polymerize gels for live cell imaging.

Once polymerized, the gels were detached from the molds and placed in a petri dish submerged with MilliQ water. Water was then substituted with phosphate buffered saline (PBS 1X) and gels were left overnight to reach the swelling equilibrium before cell seeding.

### DPAA softening

To induce controlled softening of DPAA hydrogels, glutathione (GSH, Sigma Aldrich no. PHR1359) was used as reducing agent. GSH was dissolved in Minimum Essential Medium (MEM, Thermo Fisher Scientific no. 31095029) supplemented with 10 wt% fetal bovine serum (FBS, Thermo Fisher Scientific no. A5256701), 5 wt% penicillin/streptomycin and 5 wt% glutamine to reach a final concentration of 1 mM. After complete dissolution, a proper volume of GSH solution was substituted to the culture medium to start degradation.

### Hydrogels mechanical characterization

Micropipette aspiration was used to measure bulk hydrogel stiffness. In brief, hydrogel samples were prepared attached to an adhesive glass coverslip and were left in 1× PBS overnight to reach the swelling equilibrium. For the measurement, a glass capillary connected to a syringe pump and a pressure sensor, were mounted on top of an inverted microscope with the sample holder placed perpendicular to it. The gel was placed on the holder and the capillary was moved toward the surface of the gel to achieve a complete contact. A negative pressure was then applied to the sample surface through the capillary and was detected and registered using the pressure sensor. Simultaneously, an image of the gel aspirated length inside the glass pipette was taken with the inverted microscope. The analysis of the elastic modulus was done correlating the aspirated length and the pressure exerted with the Young modulus of the hydrogel through the model published previously^38^.

The characterization of the degradation dynamics was performed preparing three samples for each timepoint. The samples were subjected to the glutathione solution and the degradation was stopped washing the samples for three times with 1X PBS. The mechanical properties were evaluated on two or three spots on the surface of each sample.

Surface hydrogel stiffness was measured by an atomic force microscopy (AFM) apparatus (Park Systems’ XE-BIO). A cantilever having a square pyramid tip was used. Young’s elastic modulus was obtained using a Hertz cone model to elaborate the indentation curves. Multiple measurements were taken every 3 minutes to characterize the degradation dynamics. The measurements were repeated three times.

### Lateral and z-swelling measurement

To evaluate the hydrogel swelling along x,y and z axis, fluorescent nanoparticles containing hydrogels were prepared and analysed acquiring confocal images of the hydrogel surface. The image acquisition was performed with a Leica Stellaris 5 confocal microscope. Degradable hydrogels were positioned in an imaging petri dish and imaged with a 10X objective for 120 minutes after the addition of a 1mM glutathione solution in culture medium. Hydrogel free surface was determined as the plane containing the last, on-focus, set of fluorescent particles. Three images of the surface were acquired every 10 minutes. Lateral swelling was evaluated comparing the relative positions of arbitrarily selected particles of the surface. Z swelling was calculated as percentual swelling relative to the initial hydrogel thickness.

### Cell culture and cell seeding

WI38 and transfected WI38 were cultured in Minimum Essential Medium (MEM) supplemented with 10 wt% fetal bovine serum (FBS), 5 wt% penicillin/streptomycin and 5 wt% glutamine. Cells were seeded on degradable hydrogels at a concentration of 2500 cells/cm^2^ for both fixed-cells analysis and live cell imaging. Fixing or live imaging were performed 24h after seeding.

### Live imaging

Cells seeded on a degradable hydrogel specifically synthetized for live imaging and polymerized on thin #0 glass substrates (90-110 μm) were treated for 60 minutes with SPY650-FastAct™ probe according to the manufacturer’s instructions. Fifteen minutes before imaging, nuclei were counterstained with a Hoechst diluted solution (1:12500 HOECHST 33342 solution in 1X PBS). Samples were finally washed twice with fresh medium, mounted on a metallic ring (Attofluor cell chamber) and filled with 400 μL of probe containing-culture medium. Live imaging was performed on a Leica Stellaris 5 confocal microscope equipped with Okolab cage incubator and a 63X oil immersion objective. Live data mode was used to establish an automated imaging protocol consisting of three steps iteratively repeated until completion of the imaging. A first step consisting of an auto-focus required to take into account the z-shifting of the gel surface during degradation is followed by a z-stack imaging of the cell. A 20 μm stack is acquired in 26 planes (every 0.8 μm). The third and final step is a pause step of 30 seconds.

Once target cell has been defined, the hydrogel softening is initiated substituting culture medium with a 1mM solution of glutathione in fresh medium. Raw files were then analyzed with Fiji software.

### Production of lentiviral particles and infection of target cells

For lentiviral particle production HEK293T were seeded at 40% confluency in 10 cm dish (Corning) and transfected with 10 μg of lentiviral plasmid containing vinculin-venus encoding gene, 2.5 μg pMD2.G (Addgene #12259) and 7.5 μg pPAX2 (Addgene#12260) by using Trans-IT X2 Dynamic Delivery system (Mirus Bio) according to manufacturer’s instructions. 48h post-transfection, supernatant was collected, filtered through 0.45 μm filter (Sarstedt) and stored at −20°C or freshly used on cells.

For in-vitro vinculin overexpression studies in WI-38 fibroblasts, cells were seeded at 30% confluency and transduced with virus diluted 1:4 in culture medium in presence of polybrene (Sigma) 8 μg/ml, for 24 hours.

The pLL3.7-CMV-Vinculin-Venus vector was generated by subcloning Vinculin-Venus (Addgene no. 27300) into the pLL3.7-CMV vector (Addgene no. 11795) by means of NheI/XbaI enzymatic digestion.

### Immunofluorescence

Immunofluorescence was performed on 4% PFA-fixed cells using anti-YAP/TAZ (Santa Cruz Biotechnology no. sc-101199 1:100), anti-vinculin (Invitrogen no. MA5-11690 1:100), anti-paxillin (abcam ab32084 1:50), anti-α5β1 (Novusbio no. NBP-252680 1:100) and anti-αvβ3 (Sigma-Aldrich MAB1976 1:100) as primary antibodies. F-actin was stained with Alexa Fluor 568 Phalloidin (Thermo Fisher Scientific no. A12380 1:100). Goat anti-mouse IgG1 Alexa fluor 647 (Thermo Fisher Scientific no. A21240 1:100) Goat anti mouse IgG2a Alexa fluor 488 (Thermo Fisher Scientific no. A21131 1:100), Goat anti-rabbit Alexa fluor 647 (Thermo Fisher Scientific no. A21245 1:100) were used as secondary antibodies. Samples were counterstained with Hoechst 33342 dye (Thermo Fisher Scientific no. 62249 1:1000) to label cell nuclei. Before imaging Confocal images were acquired with a Leica Stellaris 5 and analyzed using Fiji. The YAP/TAZ nuclear to cytosolic ratio was calculated creating a pipeline in CellProfiler software. The CellProfiler sequence was built to obtain a mask of the Nuclear Projected Area (NPA) analyzing the nuclear signal, a mask of the cellular shape based on phalloidin signal, and a mask of the cytosolic area obtained subtracting the NPA to the cell projected area. The Nuclear to Cytoplasmic ratio (N/C) was then calculated by the software on the YAP/TAZ channel as the ratio between the mean signal intensity on the nuclear mask and the mean signal intensity on the cytosolic area.

### Adhesions measurements

Adhesions length was evaluated building a custom Cellprofiler pipeline. This includes a non-local means noise reduction, a thresholding step and a segmentation step based on watershed algorithm. The pipeline than automatically calculates the major and minor axis of the detected objects. A minimum of three different ROIs for each cell have been analyzed for the peripheral adhesions while the area described by the nuclear outline was used as a mask for the analysis of the subnuclear adhesions. Data obtained were analyzed with Graphpad Prism

### Statistical analysis

Values reported are the means and standard deviations. Statistically significative differences are evaluated with a one-way ANOVA with Tukey’s multiple comparison test using GraphPad Prism 9 and considering a confidence interval of 95%.

## Supporting information

Movie 1 - F-actin live imaging

Movie 2 - Vinculin live imaging

## Acknowledgements

We thank colleagues sharing their plasmids on Addgene, and all members of the Brusatin and Piccolo groups for critical reading of the manuscript and prof. Monica Giomo for AFM measurements.

## References

1 Panciera, T., Azzolin, L., Cordenonsi, M. & Piccolo, S. Mechanobiology of YAP and TAZ in physiology and disease. Nat Rev Mol Cell Bio 18, 758–770 (2017). 10.1038/nrm.2017.87

2 Brusatin, G., Panciera, T., Gandin, A., Citron, A. & Piccolo, S. Biomaterials and engineered microenvironments to control YAP/TAZ-dependent cell behaviour. Nat Mater 17, 1063–1075 (2018). 10.1038/s41563-018-0180-8

3 Eyckmans, J., Boudou, T., Yu, X. & Chen, C. S. A hitchhiker’s guide to mechanobiology. Dev Cell 21, 35–47 (2011). 10.1016/j.devcel.2011.06.015

4 Humphrey, J. D., Dufresne, E. R. & Schwartz, M. A. Mechanotransduction and extracellular matrix homeostasis. Nat Rev Mol Cell Bio 15, 802–812 (2014). 10.1038/nrm3896

5 Elosegui-Artola, A. et al. Force Triggers YAP Nuclear Entry by Regulating Transport across Nuclear Pores. Cell 171, 1397–1410 (2017). 10.1016/j.cell.2017.10.008

6 Chen, C. S., Mrksich, M., Huang, S., Whitesides, G. M. & Ingber, D. E. Geometric control of cell life and death. Science 276, 1425–1428 (1997). DOI 10.1126/science.276.5317.1425

7 Engler, A. J., Sen, S., Sweeney, H. L. & Discher, D. E. Matrix elasticity directs stem cell lineage specification. Cell 126, 677–689 (2006). 10.1016/j.cell.2006.06.044

8 Dupont, S. et al. Role of YAP/TAZ in mechanotransduction. Nature 474, 179–U212 (2011). 10.1038/nature10137

9 Elosegui-Artola, A. et al. Rigidity sensing and adaptation through regulation of integrin types. Nature Materials 13, 631–637 (2014). 10.1038/Nmat3960

10 Gandin, A. et al. Broadly Applicable Hydrogel Fabrication Procedures Guided by YAP/TAZ-Activity Reveal Stiffness, Adhesiveness, and Nuclear Projected Area as Checkpoints for Mechanosensing. Advanced Healthcare Materials 11 (2022). 10.1002/adhm.202102276

11 Quinlan, A. M. T. & Billiar, K. L. Investigating the role of substrate stiffness in the persistence of valvular interstitial cell activation. J Biomed Mater Res A 100a, 2474–2482 (2012). 10.1002/jbm.a.34162

12 Yui, S. et al. YAP/TAZ-Dependent Reprogramming of Colonic Epithelium Links ECM Remodeling to Tissue Regeneration. Cell Stem Cell 22, 35–49 (2018). 10.1016/j.stem.2017.11.001

13 Panciera, T. et al. Reprogramming normal cells into tumour precursors requires ECM stiffness and oncogene-mediated changes of cell mechanical properties. Nat Mater 19, 797–806 (2020). 10.1038/s41563-020-0615-x

14 Fan, W. G. et al. Matrix viscoelasticity promotes liver cancer progression in the pre-cirrhotic liver. Nature 626 (2024). 10.1038/s41586-023-06991-9

15 Mabry, K. M., Lawrence, R. L. & Anseth, K. S. Dynamic stiffening of poly(ethylene glycol)-based hydrogels to direct valvular interstitial cell phenotype in a three-dimensional environment. Biomaterials 49, 47–56 (2015). 10.1016/j.biomaterials.2015.01.047

16 Ondeck, M. G. & Engler, A. J. Mechanical Characterization of a Dynamic and Tunable Methacrylated Hyaluronic Acid Hydrogel. J Biomech Eng-T Asme 138 (2016). 10.1115/1.4032429

17 Guvendiren, M. & Burdick, J. A. Stiffening hydrogels to probe short- and long-term cellular responses to dynamic mechanics. Nat Commun 3 (2012). 10.1038/ncomms1792

18 Chaudhuri, O. et al. Hydrogels with tunable stress relaxation regulate stem cell fate and activity. Nature Materials 15, 326–334 (2016). 10.1038/Nmat4489

19 Loebel, C., Mauck, R. L. & Burdick, J. A. Local nascent protein deposition and remodelling guide mesenchymal stromal cell mechanosensing and fate in three-dimensional hydrogels. Nature Materials 18, 883–891 (2019). 10.1038/s41563-019-0307-6

20 Roca-Cusachs, P., Iskratsch, T. & Sheetz, M. P. Finding the weakest link - exploring integrin-mediated mechanical molecular pathways. J Cell Sci 125, 3025–3038 (2012). 10.1242/jcs.095794

21 Kloxin, A. M., Benton, J. A. & Anseth, K. S. elasticity modulation with dynamic substrates to direct cell phenotype. Biomaterials 31, 1–8 (2010). 10.1016/j.biomaterials.2009.09.025

22 Yang, C., Tibbitt, M. W., Basta, L. & Anseth, K. S. Mechanical memory and dosing influence stem cell fate. Nature Materials 13, 645–652 (2014). 10.1038/Nmat3889

23 Wang, H., Haeger, S. M., Kloxin, A. M., Leinwand, L. A. & Anseth, K. S. Redirecting Valvular Myofibroblasts into Dormant Fibroblasts through Light-mediated Reduction in Substrate Modulus. Plos One 7 (2012). 10.1371/journal.pone.0039969

24 Tse, J. R. & Engler, A. J. Preparation of hydrogel substrates with tunable mechanical properties. Curr Protoc Cell Biol Chapter 10, Unit 10 16 (2010). 10.1002/0471143030.cb1016s47

25 Prasetyanto, E. A. et al. Breakable Hybrid Organosilica Nanocapsules for Protein Delivery. Angew Chem Int Edit 55, 3323–3327 (2016). 10.1002/anie.201508288

26 Moghaddam, S. P. H., Yazdimamaghani, M. & Ghandehari, H. Glutathione-sensitive hollow mesoporous silica nanoparticles for controlled drug delivery. Journal of Controlled Release 282, 62–75 (2018). 10.1016/j.jconrel.2018.04.032

27 Gyarmati, B., Nemethy, A. & Szilagyi, A. Reversible disulphide formation in polymer networks: A versatile functional group from synthesis to applications. Eur Polym J 49, 1268–1286 (2013). 10.1016/j.eurpolymj.2013.03.001

28 Cui, C. Y., Li, B. & Su, X. C. Real-Time Monitoring of the Level and Activity of Intracellular Glutathione in Live Cells at Atomic Resolution by (19)F-NMR. ACS Cent Sci 9, 1623–1632 (2023). 10.1021/acscentsci.3c00385

29 Cnubben, N. H., Rietjens, I. M., Wortelboer, H., Van Zanden, J. & Van Bladeren, P. J. The interplay of glutathione-related processes in antioxidant defense. Environ Toxicol Pharmacol 10, 141–152 (2001). 10.1016/s1382-6689(01)00077-1

30 Wilson, M. J., Liliensiek, S. J., Murphy, C. J., Murphy, W. L. & Nealey, P. F. Hydrogels with well-defined peptide-hydrogel spacing and concentration: impact on epithelial cell behavior. Soft Matter 8, 390–398 (2012). 10.1039/c1sm06589k

31 Vining, K. H. & Mooney, D. J. Mechanical forces direct stem cell behaviour in development and regeneration. Nat Rev Mol Cell Bio 18, 728–742 (2017). 10.1038/nrm.2017.108

32 Kechagia, J. Z., Ivaska, J. & Roca-Cusachs, P. Integrins as biomechanical sensors of the microenvironment. Nat Rev Mol Cell Bio 20, 457–473 (2019). 10.1038/s41580-019-0134-2

33 Kim, D. H., Hah, J. & Wirtz, D. Mechanics of the Cell Nucleus. Adv Exp Med Biol 1092, 41–55 (2018). 10.1007/978-3-319-95294-9_3

34 Kirby, T. J. & Lammerding, J. Emerging views of the nucleus as a cellular mechanosensor. Nat Cell Biol 20, 373–381 (2018). 10.1038/s41556-018-0038-y

35 Shiu, J. Y., Aires, L., Lin, Z. & Vogel, V. Nanopillar force measurements reveal actin-cap-mediated YAP mechanotransduction. Nat Cell Biol 20, 262–271 (2018). 10.1038/s41556-017-0030-y

36 Elosegui-Artola, A. et al. Mechanical regulation of a molecular clutch defines force transmission and transduction in response to matrix rigidity. Nat Cell Biol 18, 540–548 (2016). 10.1038/ncb3336

37 Oria, R. et al. Force loading explains spatial sensing of ligands by cells. Nature 552, 219–224 (2017). 10.1038/nature24662

38 Gandin, A. et al. Simple yet effective methods to probe hydrogel stiffness for mechanobiology. Sci Rep-Uk 11 (2021). 10.1038/s41598-021-01036-5

