## Supplemental Figures for "YAP/TAZ mechanotransduction is a fast cellular response to subnuclear adhesions revealed by mechanically dynamic substrates"

**Table S1:** Chemical compounds used for the hydrogel synthesis and functionalization.

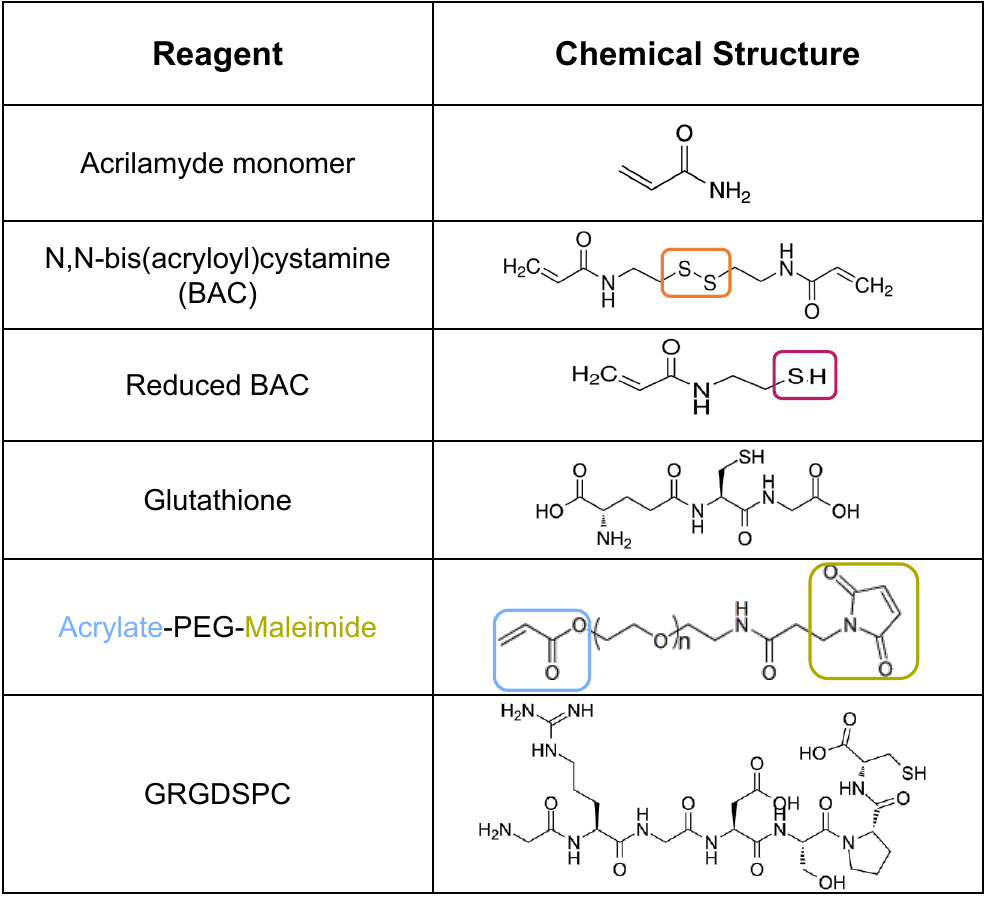

**Table S2:** Composition of different DPAA formulations prepared with 3mM of RGD. The initial modulus (E_in_, measured after 1 day swelling) and the final modulus (E_1h_, measured after 1h degradation, with 1mM GSH in PBS solution) are measured by micropipette aspiration (columns 3 and 4 respectively).

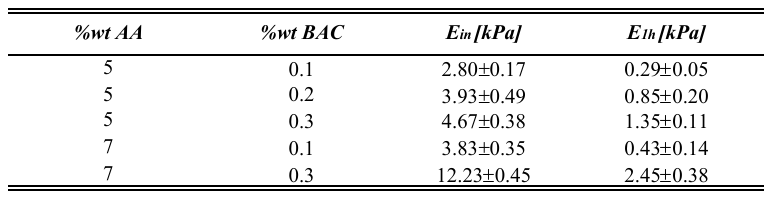

**Table S3:** Elastic modulus of degradable gel used for the seeding experiments (7%AA, 0.3%BAC, 3 mM of RGD), after different degradation times with GSH 0.5 or 1mM. The moduli are measured by the micropipette aspiration method.

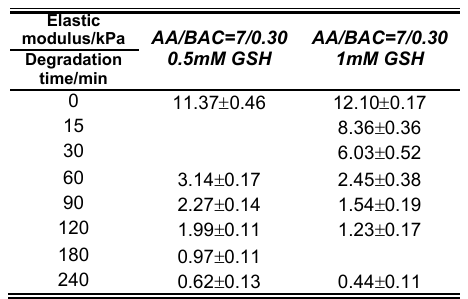

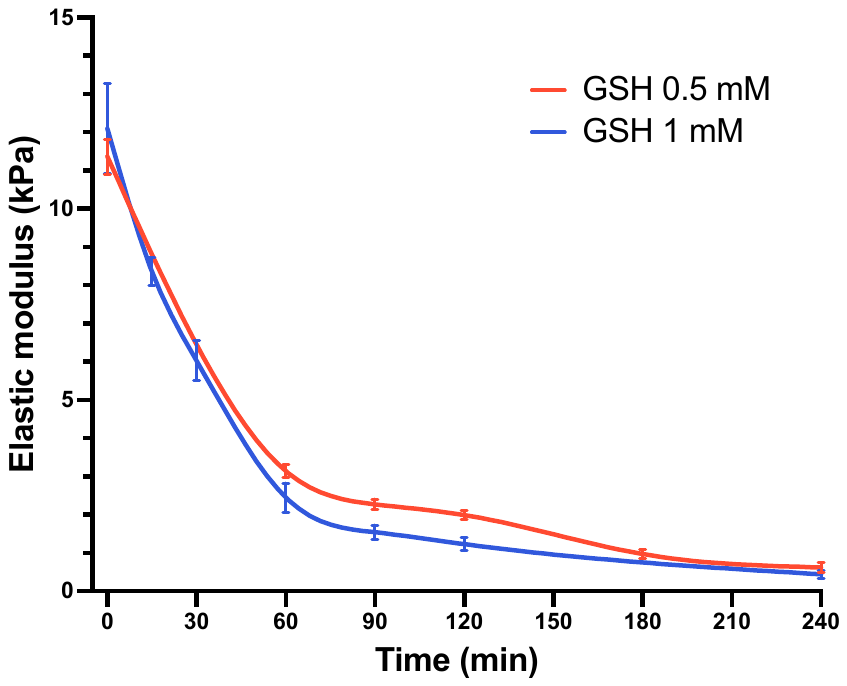

**Figure S1**: Bulk stiffnesses measured at different time points during gel softening with an initial stiffness of about 12kPa with 0.5 or 1mM GSH for 4h. The degradation has been followed until a plateau value of stiffness is reached.

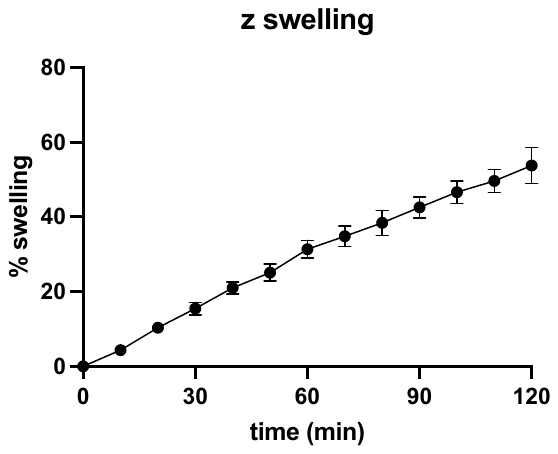

**Figure S2**: Swelling behavior along the z axis (thickness) measured during gel softening (GSH 1 mM). Measurements were acquired recording the position of the gel surface through confocal imaging of fluorescent nanoparticles embedded in the hydrogel

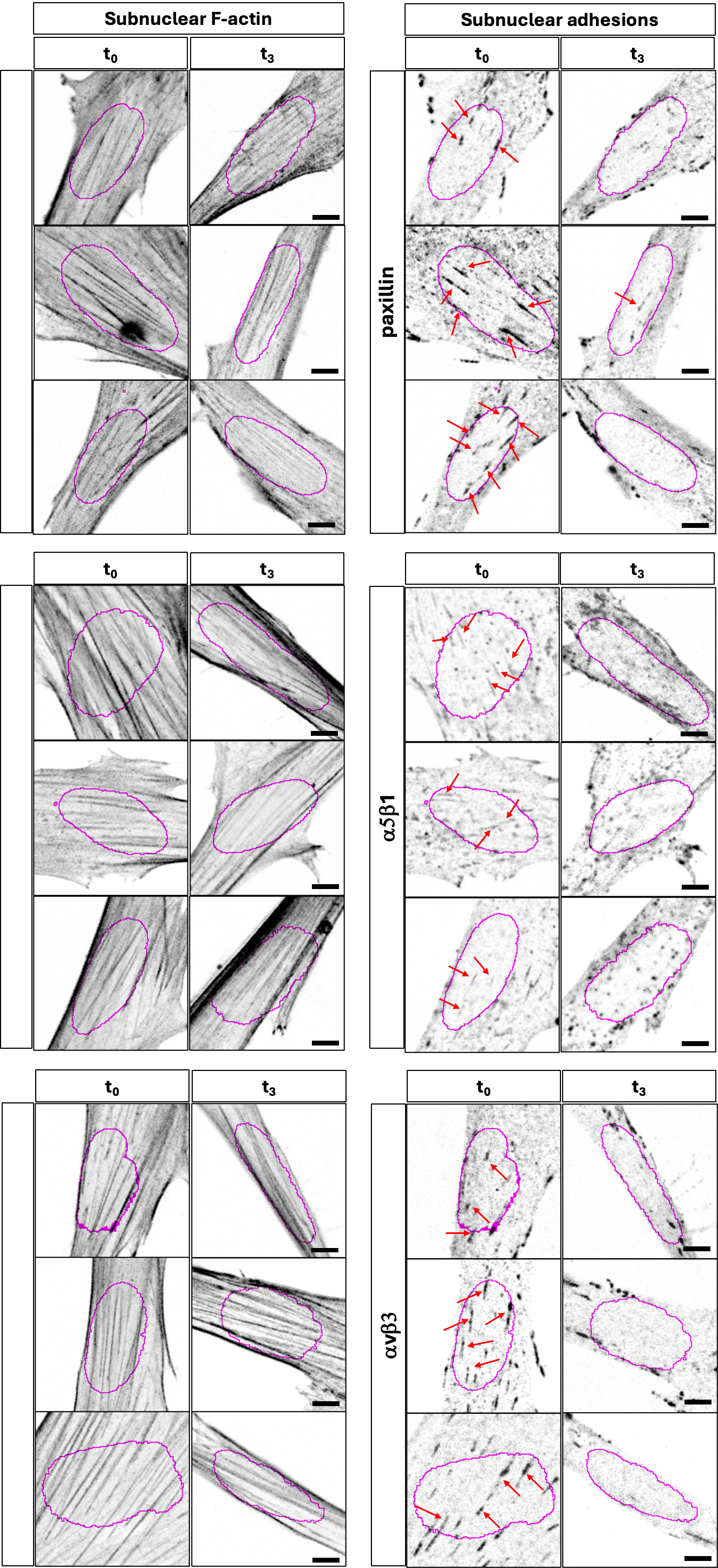

**Figure S3**: Representative immunofluorescence images of cell on DPAA gel with an initial stiffness of about 12 kPa (t0) and during softening (t3). Subnuclear basal actin and cellular adhesions (paxillin, a_5_b_1_ or a_v_b_3_) were imaged in the same cell through confocal microscopy. Magenta circles represent the outlines of the nuclear projected area. Red arrows indicate the main subnuclear adhesions. Scale bar 5 μm
